# Weak evaporative cooling capacity and body size shape thermal limits in tropical montane forest birds

**DOI:** 10.1101/2025.11.11.687810

**Authors:** Cesare Pacioni, Ran Xu, Beate Apfelbeck, Frederick Verbruggen, Peter Njoroge, Mwangi Githiru, Diederik Strubbe, Luc Lens

## Abstract

Climate warming and forest fragmentation threaten tropical biodiversity by altering microclimates and narrowing the range of temperatures within which species can maintain optimal physiological performance. We investigated the thermoregulatory capacities of five forest-dependent bird species from the Taita Hills (Kenya), a montane biodiversity hotspot experiencing severe habitat loss. Using respirometry, we measured resting metabolic rate (RMR) across a range of temperatures to determine thermoneutral zone (TNZ) limits, quantified evaporative water loss (EWL), evaporative heat loss (EHL), and metabolic heat production (MHP) as proxies of cooling capacity, and estimated heat tolerance limits (HTLs). Contrary to expectations of a classical TNZ pattern, RMR–temperature relationships were predominantly V-shaped, suggesting the absence, or marked narrowness, of a TNZ. In contrast, HTL clearly increased with body mass, with larger species tolerating higher temperatures. Evaporative cooling efficiency remained weak across all species (EHL/MHP < 1), indicating limited capacity to dissipate metabolic heat. Compared with global data, the Taita Hills species exhibited low critical temperatures and narrow thermal ranges, consistent with specialisation to the stable microclimates of tropical montane forests. Our findings suggest that small-bodied, forest-dependent tropical birds function within narrow thermal margins, which may make them especially vulnerable to rising temperatures and the microclimatic changes associated with climate change and forest fragmentation.

## 1. Introduction

The Earth’s climate and ecosystems are experiencing rapid, unprecedented changes due to human activities (IPCC, 2023). With rising average temperatures and an increased frequency of extreme heatwaves, understanding how organisms will adapt to these shifts has become one of the most urgent challenges in ecology and conservation biology (Schwenk et al., 2009; Stillman, 2019). In this context, forest fragmentation acts as an amplifier of these effects, with 70% of the world’s remaining forests located within 1 km of their edges (Haddad et al., 2015), often surrounded by landscapes with low structural complexity (Linke, 2007). Such altered land covers can increase solar radiation exposure and modify microclimatic conditions within forest patches (Tuff et al., 2016), thereby exacerbating the physiological stress imposed by climate warming. Projections suggest that fragmentation will worsen over the next five decades, particularly in tropical regions, through reduced patch sizes, greater distances between patches, and forest loss (Alroy, 2017; Barlow et al., 2016; Taubert et al., 2018). Although habitat fragmentation has been widely studied for its impacts on species and communities, the links between altered microclimates and species’ thermal physiology remain poorly understood (Monge et al., 2023; Tuff et al., 2016). This gap is concerning, as the combined effects of global warming and forest fragmentation represent one of the most pressing threats to biodiversity and ecosystem services (IPBES, 2019).

In this context, understanding species’ physiological limits is essential to assess their vulnerability to changing thermal environments (Bozinovic & Pörtner, 2015). An important physiological parameter that governs how organisms cope with thermal stress is the thermoneutral zone (TNZ). This refers to the range of ambient temperatures within which an endotherm can maintain a stable body temperature without increasing its resting metabolic rate (RMR) above the basal metabolic rate (BMR) required to sustain basic life functions (McNab, 2012; Ruuskanen et al., 2021). When ambient temperatures fall outside the TNZ, endotherms must spend additional energy to maintain internal temperature homeostasis (McNab, 2012). For example, higher temperatures can expose species to thermal stress that may constrain activity, or breeding performance, and ultimately, under extreme conditions such as during heat waves, exceed physiological tolerance thresholds and cause mortality (Cabello-Vergel et al., 2022; Pollock et al., 2021). To cope with the increased heat load, endotherms primarily rely on evaporative water loss (EWL) to dissipate excess heat (McNab, 2012), albeit with the potential risk of dehydration if species are unable to replenish their water reserves (Mckechnie & Swanson, 2010; Swanson, 2010). Organisms, however, are not limited to physiological mechanisms in coping with thermal stress. Morphological traits, such as plumage coloration, feather structure and density, and body size, can modulate heat absorption and dissipation (Osváth et al., 2018; Rogalla et al., 2021; Wolf & Walsberg, 1996). Likewise, behavioural strategies such as shade-seeking, altering activity patterns, or using thermally buffered microhabitats can substantially reduce heat exposure (Carroll et al., 2015; Scheffers et al., 2014). These buffering strategies often act in concert with physiological responses, and their relative importance may vary across species and environments. However, in fragmented tropical forests, the availability of suitable microhabitats or structural cover is often reduced, which may limit the effectiveness of such strategies and exacerbate vulnerability to heat stress.

Among endotherms, diurnally-active, small birds are particularly susceptible to heat stress due to their smaller body water pool and energy stores, combined with high surface area-to-volume ratios (Albright et al., 2017). Studies have shown that small birds can lose more than 5% of their body mass per hour through evaporative cooling, even while inactive and in fully shaded microhabitats (McKechnie & Wolf, 2009; Wolf & Walsberg, 1996). This high rate of mass loss is linked to their lower evaporative cooling efficiency (defined as the quotient between evaporative heat loss, EHL, and metabolic heat production, MHP; Cabello-Vergel et al., 2022), as smaller species often lack the effective heat dissipation mechanisms found in larger-bodied birds (McKechnie et al., 2021a). Small birds therefore often have lower heat tolerance limits (HTL), which are the highest air temperatures they can endure body temperature rises uncontrollably and evaporative cooling fails to prevent lethal overheating (Conradie et al., 2020; McKechnie et al., 2021b). Body size may further modulate thermoregulatory performance, as larger birds typically have greater heat storage capacity and thermal inertia, which could allow them to maintain stable body temperature across a wider range of ambient temperatures (McNab, 2002). Moreover, small bird species living in tropical areas may already be near their maximum thermal tolerance (Araújo et al., 2013) and climate change could push them beyond their limits, especially in combination with forest fragmentation.

This vulnerability is particularly evident for habitat-dependent species (Keinath et al., 2017). Many tropical forest-dependent birds, especially those inhabiting montane forests, are habitat and thermal specialists, adapted to the stable, humid conditions of forest interiors and typically tolerating only a narrow range of temperatures (Polato et al., 2018). This suggests that even modest increases in ambient temperature or microclimatic variation could impose substantial thermoregulatory costs. Adaptation to the stable conditions of montane forests may have resulted in narrower TNZs, lower critical temperatures, and reduced efficiency of evaporative heat loss relative to metabolic heat production (EHL/MHP) compared to species from hotter or more variable environments. Consequently, tropical montane forest birds are expected to exhibit a limited capacity for heat dissipation and overall weaker thermoregulatory performance. However, most studies on avian thermal physiology have focused on desert and arid-zone species, leaving a substantial gap in our understanding of how forest-dependent tropical birds cope with heat stress. In particular, basic physiological data on their thermal limits, evaporative cooling efficiency, and heat tolerance thresholds are lacking, making it difficult to predict how species will respond to warming, especially in fragmented forest landscapes (Monge et al., 2023).

Here, we assessed the thermoregulatory capacity of free-living tropical birds inhabiting a fragmented montane forest by i) measuring resting metabolic rate (RMR) across a range of air temperatures to determine the limits of the thermoneutral zone (TNZ), ii) quantifying evaporative water loss (EWL), evaporative heat loss (EHL), and metabolic heat production (MHP) as indicators of cooling capacity, and iii) estimating heat tolerance limits (HTL). The study focused on five small-to medium-sized, forest-dependent species from the Taita Hills (Kenya), part of the Eastern Afromontane biodiversity hotspot, where severe forest loss has left remaining patches isolated and threatened, despite the region’s high species richness and endemism. In particular, we studied: the olive sunbird (*Cyanomitra olivacea*), olive-headed greenbul (*Andropadus milanjensis*), Taita white-eye (*Zosterops silvanus*), white-starred robin (*Pogonocichla stellata*), and yellow-throated woodland warbler (*Phylloscopus ruficapilla*). These species, ranging from ∼7 to ∼43 g in body mass, are highly dependent on forest habitats and persist across all forest fragments in the Taita landscape (Mulwa et al., 2021), making them suitable subjects for testing interspecific variation in thermoregulatory performance.

We hypothesized that (1) larger-bodied bird species would exhibit broader TNZs than smaller species; (2) larger-bodied species would also cope more effectively with heat, showing (2a) higher HTLs and (2b) greater evaporative cooling capacity compared to smaller species (McKechnie et al., 2021a); and (3) all study species, being tropical montane forest specialists adapted to stable, humid, and moderate environments, would display weak evaporative cooling efficiency (EHL/MHP < 1). To place our findings in a broader biogeographic context, we compared the thermoregulatory parameters of our study species with the global dataset compiled by Khaliq et al. (2015). Given that the Taita Hills represent a montane tropical environment characterized by relatively stable and moderate temperatures, cooler than surrounding lowlands but not cold by temperate standards, we expected that (4) forest-dependent species would exhibit thermal traits reflecting adaptation to the stable climatic conditions of montane forests, specifically (4a) narrow TNZs and (4b) both lower and upper critical temperatures occurring at relatively low ambient temperatures, consistent with specialization to these mild montane conditions rather than to hot or highly variable environments.

## 2. Material and methods

The forest fragments of the Taita Hills (Kenya) are part of the Eastern Afromontane biodiversity hotspot, considered one of the top 10 biodiversity hotspots in the world (Lovett & Wasser, 2008). They mark the northernmost extent of the Eastern Arc Mountains, a mountain range that stretches from south-eastern Kenya to southern Tanzania and is known for its rich biodiversity, high endemism, and considerable threat levels (Burgess et al., 2007). It is estimated that only 30% of the region’s original forest cover remains (Myers et al., 2000; Newmark, 2002). In particular, the Taita Hill forests are among the most severely impacted and endangered within the Eastern Arc mountain range (Teucher et al., 2020). Approximately 90% of the original indigenous forest cover has been lost to agricultural activities, while selective logging for firewood has further degraded the interior of the remaining fragments (Lovett & Wasser, 2008; Newmark, 2002; Pellikka et al., 2009). Despite their small size, the 12 remaining fragments of native forest play a critical role in global conservation efforts by supporting rare and endemic plants and animals (Pellikka et al., 2013). However, these fragments face immediate conservation challenges, including increasing isolation, soil erosion, and adverse hydrological impacts, coupled with the challenges posed by climate change (Pellikka et al., 2013; Terschanski et al., 2024). For these reasons, they offer a unique setting for studying the thermal physiology of tropical species.

The study was carried out in 2025 during the non-breeding season. Birds were attracted using song playback and captured using mist nets in one of the Taita Hills’ (03°20′S, 38°15′E) forest fragments, Ngangao. Birds showing a brood patch indicative of breeding activity were promptly released. The remaining individuals were ringed, transferred to cloth bags, and quickly transported to a nearby facility, where they were housed in perch-equipped cages and provided with water ad libitum. Nocturnal RMR and EWL measurements started on the evening the birds were captured. The following morning, one or two individuals were kept for HTL measurements, provided with food and water until HTL measurements, and released later that day, after the measurements were completed. The other birds were housed with water provided before being released at the capture site. To minimize stress and ensure animal welfare, each individual was measured only once. In the rare case of recapture, the bird was released immediately without additional measurements. Species names were cross-referenced and standardized according to the unified global checklist provided by AviList (https://www.avilist.org; Rheindt et al., 2025). The experimental protocols were approved by the IRC-UGent Ethical Committee Animal Experimentation with reference code EC2025-024 and by the Wildlife Research and Training Institute (WRTI) with reference code WRTI/RPC/2024-470663.

### 2.1. Experimental protocol

RMR was measured at night using an open flow-through respirometry system. The RMR at different temperatures was used to quantify the TNZ of the species. The lowest RMR was considered BMR. Birds were placed in airtight plastic chambers of different volumes (1 L, 2.3 L, or 3.7 L) depending on their size and tail length, and were kept from dusk to dawn (approximately 12 hours) in a darkened, climate-controlled unit (PELT-5; Sable Systems). RMR was assessed over a range of air temperatures, from 10 to 32 °C. This temperature range was selected to encompass a broad spectrum of thermal conditions experienced by the study species in the study area (WordClim data) and thus to assess their thermoregulatory capacity. Several temperatures were tested per night (∼3 hours per temperature, following van de Ven et al., 2013). The first 30 minutes of each tested temperature were discarded from the analysis to ensure the birds were acclimatized to that temperature. Water vapor was removed from the air stream immediately downstream of the metabolic chambers using Drierite® (Lighton, 2018). A pump (PP-2-1; Sable Systems) supplied dehumidified air, which was then divided into eight separate streams and directed to a mass-flow meter (FB-8; Sable Systems). Needle valves of the mass-flow meter were adjusted to maintain an appropriate flow rate (between 400-600 mL minLJ¹, depending on bird size and chamber volume) that kept COLJ concentrations below 0.5%. Excurrent air from each chamber, as well as the baseline channel, was sampled alternately via a multiplexer (RM-8; Sable Systems) for independent measurement of each chamber. Excurrent air from the bird and the baseline channels was alternately subsampled and pulled through a Field Metabolic System (FMS-3; Sable Systems). Birds and baseline measurements were alternated in cycles, with the cycle length and measurement time for each bird depending on the number of birds in the session (on average, ∼2 individuals per night). After respirometry, birds were weighed to the nearest 0.1 g and given water for approximately one hour before being released at the capture site.

For nocturnal EWL (VHLJO) measurements, the setup was similar to that used for RMR measurements. Birds were exposed to a range of temperatures from their TNZ up to 32 °C (see above). During the EWL measurements, the birds stood on a platform 10 cm above a 1 cm layer of mineral oil, which trapped any excreta. This setup ensured that evaporative water loss was measured without contamination from waste. Temperature changes were controlled automatically within the climate-controlled unit, following the same protocol as for RMR. The water vapor was removed from the air stream downstream of the chambers using Drierite® (Lighton, 2018).

HTL measurements were conducted following a similar procedure employed in recent studies (e.g., Cabello-Vergel et al., 2022). Briefly, heat tolerance trials were carried out during the birds’ active phase, with each bird measured individually using a similar setup as for the nocturnal EWL measurements. Before the measurements, birds were given one hour of rest in a dark place with food and water after the RMR and EWL measurements from the night before. Gas exchange rates were measured using a stepped temperature profile that started within the TNZ of the species (assessed at the beginning of the study) and increased in 2 °C increments. The temperature was maintained for a minimum of 10 minutes at each target temperature, during which baseline air samples were collected for 1 minute at the beginning of each temperature step to ensure proper calibration. At each temperature, the bird’s oxygen consumption and water vapor output were constantly monitored. This stepped approach was efficient, reducing the time that the birds were confined to the chamber compared to steady-state protocols, which exposed birds to each temperature for longer periods (Whitfield et al., 2015). The trial ended when the bird exhibited signs of heat stress, such as sudden decreases in oxygen consumption or water vapor production. The maximum temperature reached reflected the bird’s critical HTL.

### 2.2. Respirometry and data analysis

The software ExpeData (Sable Systems) was used for trial recording and extraction of metabolic rate values (mL OLJ minLJ¹), including correcting for O_2_ and H_2_O drift. To estimate RMR and EWL, in ExpeData, the lowest stable portion of the curve (average of 5 min) at each temperature set point was selected and RMR and EWL were estimated using equation 9.7 and 10.9 from Lighton (2018), respectively. In these equations, incurrent %HLJO was set to zero, as water was eliminated using Drierite®. RMR was converted to MHP (W) using the conversion factor 1LJmL O_2_ = 20.083LJJ (Schmidt-Nielsen, 1997). EWL was converted to EHL (W) assuming a latent heat of vaporization of 2.406 J mg^−1^ (Tracy, 2010). The quotient between EHL and MHP (at each temperature tested) was considered a proxy of cooling capacity. An EHL/MHP value greater than 1 signifies that an individual can dissipate all its metabolic heat through evaporation.

### 2.3. Statistical analysis

All analyses were conducted in R v. 4.3.1 (R Core Team, 2023). To determine the limits of the TNZ, we modelled the relationship between ambient temperature and RMR for each species using a set of increasingly complex candidate models. A linear model was used as a null model, representing no change in RMR with temperature. A one-breakpoint model allowed for a single change in slope, corresponding to a potential lower or upper critical temperature, while a two-breakpoint model represented the expected pattern of a classic TNZ, with RMR remaining constant between lower and upper critical limits. Models were fitted using the *segmented* package (Muggeo, 2008). Although we expected a two-breakpoint relationship based on standard endotherm physiology, we included the simpler models to assess whether the data provided sufficient statistical support for the presence of one or two breakpoints and to avoid over-parameterization when evidence for a plateau was weak. TNZ width was calculated as the difference between the upper and lower breakpoints for species with two identifiable breakpoints. Model support was assessed using AIC, comparing linear, one-breakpoint, and two-breakpoint models. Models within ΔAIC ≤ 2 of the best-supported model were considered to have comparable support.

The association between TNZ width and log-transformed body mass was examined under the assumption that the two-breakpoint model provided the best representation of the RMR–temperature relationship, thereby yielding measurable lower and upper critical temperatures for each species. This analysis aimed to test whether larger-bodied species exhibited broader TNZs than smaller species. Non-parametric bootstrapping (10,000 replicates) was used to estimate the empirical distribution of the slope, generating robust 95% confidence intervals. Replicates were obtained by sampling species with replacement, fitting the regression, and extracting the slope. Non-finite slopes (<2% of iterations) were discarded. Permutation testing (10,000 iterations) further assessed the significance of the observed slope by randomly shuffling body mass across species and recalculating slopes, with the proportion of permuted slopes exceeding the observed slope providing a two-sided p-value. Because TNZ width is a species-level trait (one data point per species), we used bootstrapping and permutation tests to account for the limited scope for parametric inference. In addition to OLS regression with bootstrapping and permutation tests, we fitted a Bayesian linear regression to estimate the posterior distribution of the slope relating TNZ width to log-transformed body mass. We used weakly informative priors centred at zero with a standard deviation large enough to allow a wide range of plausible slopes, reflecting minimal prior knowledge about the expected relationship. Additionally, Cook’s distance was calculated for each species to assess the influence of individual observations on the regression slope. Cook’s distance values exceeding the conventional threshold 4/(n-2) were considered to indicate disproportionate leverage.

Heat tolerance limits (HTLs) were analysed using the same species-level approach as TNZ width. For evaporative cooling, segmented regression was also used to identify the ambient temperature above which EHL/MHP changed most rapidly. Following Cabello-Vergel et al. (2022), the slope above the upper inflection point (or, if no upper inflection existed, the only inflection point identified in the segmented regression models) was extracted to quantify each species’ capacity for evaporative heat loss at high temperatures. These slopes were then related to mean species log-transformed body mass using linear models.

To place our findings in a broader biogeographic context (prediction 4), the TNZ parameters of the study species from the Taita Hills were compared with a global dataset compiled by Khaliq et al. (2015), restricted to bird species with body masses ≤ 40 g. Lower (LCT) and upper (UCT) critical temperatures, as well as TNZ width, were extracted for all species and for tropical/subtropical species (latitude ±30°). The empirical cumulative distribution function (ECDF) was used to determine the percentile positions of the Taita Hills species for each variable relative to these global distributions. Density distributions of LCT, UCT, and TNZ width were plotted for both the global and tropical subsets. To contextualize the elevation of the Taita Hills study site (∼1,800LJm), occurrence points of the tropical small-bird subset were compiled, and elevations were retrieved via the Open-Elevation API; these ranged from near sea level to approximately 1,514 m, placing the study site above the maximum elevation represented in the subset.

For all analyses, three body mass measurements (pre-, post-, and mean values) were tested. Since no significant differences in outcome were observed, post-measurements are reported here, as they best represent the birds’ actual body mass after an overnight fast, without recent food intake, and are therefore closest to their baseline physiological mass. Outliers were checked but none were detected. Residual normality was verified using the Shapiro–Wilk test (W > 0.9). All traits were log-transformed prior to analysis to meet model assumptions. Statistical significance was set at p ≤ 0.05. Additional details and R scripts are provided in the Supplementary File (RMarkdown HTML).

Given the small number of species in this study, all species-level analyses were exploratory in nature, aimed at identifying potential patterns in thermoregulatory traits and generating hypotheses for future work. Moreover, we conducted additional exploratory analyses by estimating Pagel’s λ and fitting preliminary Phylogenetic Generalized Least Squares (PGLS) models. These analyses did not change the direction, magnitude, or interpretation of the main results, and therefore did not alter our conclusions. While reassuring, these exploratory outcomes must be interpreted with caution, as estimating parameters such as λ or fitting PGLS models can be unreliable with so few species, since robust phylogenetic inference may require larger sample sizes relative to the number of predictors (Marcondes, 2019; Mundry, 2014; Pearse et al., 2025). In addition, λ estimates in very small or incompletely sampled clades (e.g., the Taita white-eye, which is absent from many commonly used phylogenies) are highly uncertain and particularly sensitive to violations of assumptions, such as the requirement of random species sampling (Marcondes, 2019). For this reason, we focused on observed interspecific differences rather than phylogenetically generalized effects, and we acknowledge that formal phylogenetic correction is not feasible under the present conditions.

## 3. Results

Across all five study species, the relationship between RMR and ambient temperature exhibited a broadly V-shaped form (Figure 1). Statistically, a two-breakpoint model was selected for two species, the olive sunbird and the white-starred robin, while a one-breakpoint (i.e., V-shaped) model received strongest support for the remaining three (Table 1). However, even for the two species with apparent support for a two-breakpoint relationship, the estimated breakpoints did not define a physiologically credible TNZ: one breakpoint broadly coincided with the temperature of minimum RMR, whereas the second occurred at much higher temperatures where metabolism had already increased. The temperatures at which minimum RMR was achieved were broadly similar among species, averaging around 25 °C (range = 23.7–26.2 °C). Post-hoc pairwise comparisons of these minima, based on *Z*-tests of the estimated values and their standard errors and corrected for multiple testing, indicated limited interspecific differences, with only the olive-headed greenbul exhibiting a significantly higher minimum than several other species (Table S1 – Supplementary material).

**Figure 1.**
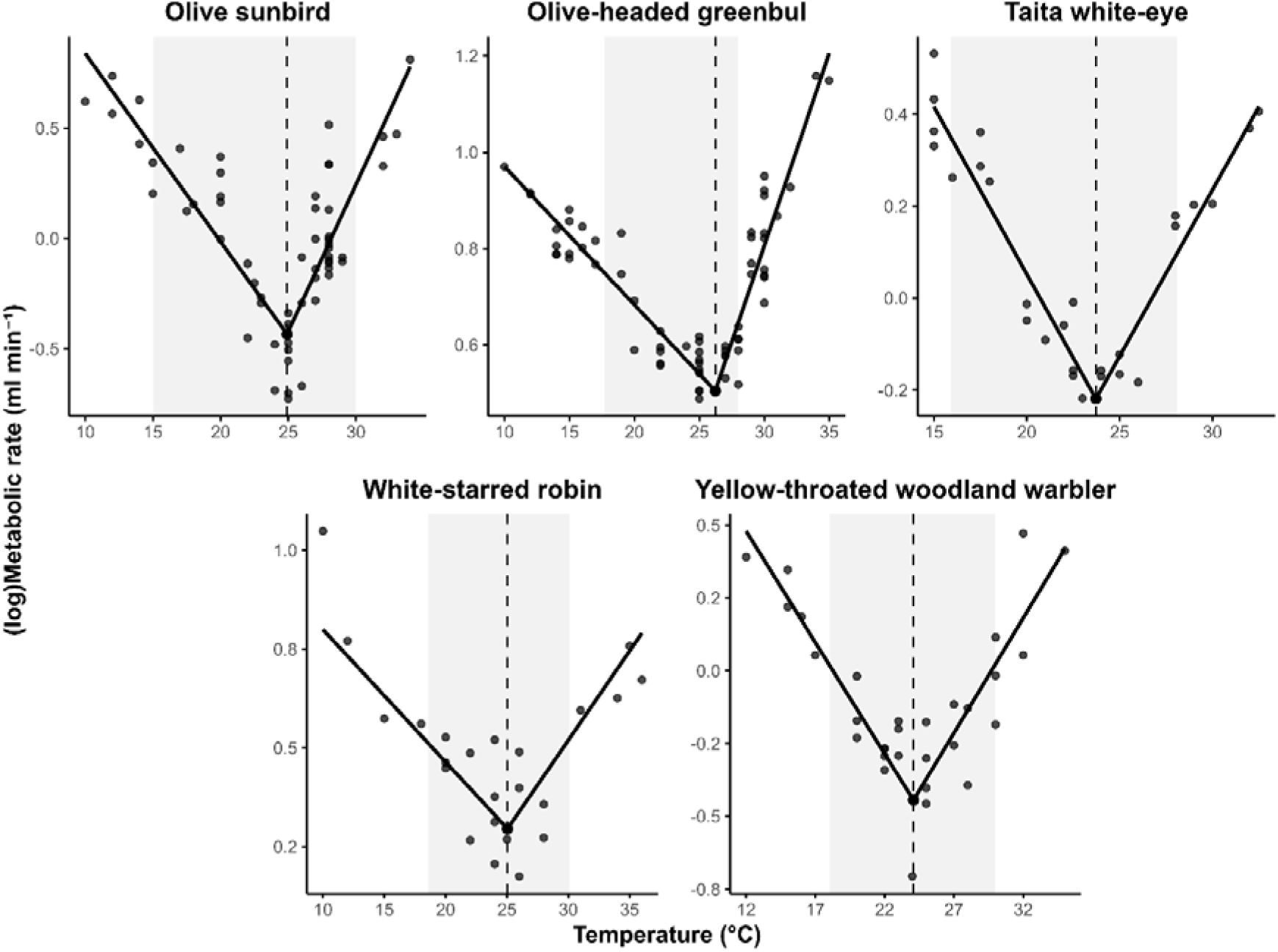
Relationship between log-transformed metabolic rate and ambient temperature for five tropical montane forest bird species. Each panel shows one species. with metabolic rate values plotted against air temperature (°C). Solid lines depict segmented regression fits describing the decline and subsequent increase in metabolic rate around a single estimated breakpoint (dashed vertical line). The shaded grey bands indicate species-specific temperature ranges corresponding to the thermoneutral zone (TNZ) as estimated from two-breakpoint models. though these were not considered ecologically realistic.

**Table 1.** Results from segmented regression analyses of log-transformed rest metabolic rate (RMR) versus ambient temperature. Models include linear, 1-breakpoint, and 2-breakpoint fits. AIC values indicate model fit. Breakpoints (°C) are estimated inflection points in the RMR–temperature relationship, with ± representing the standard deviation of each estimate, where applicable.

| Species | Model | AIC | Breakpoint #1 °C | Breakpoint #2 °C |
| --- | --- | --- | --- | --- |
| Olive sunbird | Linear | 54.61950994 |  |  |
|  | 1-breakpoint | -17.80130119 | 24.91±0.43 |  |
|  | 2-breakpoint | -23.16076248 | 20.00±2.29 | 24.00±0.43 |
| Olive-headed greenbul | Linear | -48.51127122 |  |  |
|  | 1-breakpoint | -173.5238261 | 26.23±0.25 |  |
|  | 2-breakpoint | -170.4411686 | 17.16±4.98 | 26.14±0.28 |
| Taita white-eye | Linear | 2.298014784 |  |  |
|  | 1-breakpoint | -57.32398012 | 23.72±0.37 |  |
|  | 2-breakpoint | -54.11690298 | 21.00±2.36 | 24.29±0.60 |
| Yellow-throated woodland warbler | Linear | 14.08711883 |  |  |
|  | 1-breakpoint | -28.59421295 | 24.08±0.54 |  |
|  | 2-breakpoint | -25.62423198 | 24.00±1.26 | 26.77±7.68 |
| White starred-robin | Linear | -6.20994 |  |  |
|  | 1-breakpoint | -44.0526 | 25.04±0.54 |  |
|  | 2-breakpoint | -53.0186 | 12.92±1.26 | 27.01±7.68 |

Because the metabolic curves of all study species were predominantly V-shaped, lower and upper critical temperatures effectively coincided, leaving no measurable TNZ width to analyse. Consequently, the relationship between TNZ width and body mass (prediction 1) could not be meaningfully tested. In line with our prediction 2a, species-level HTL increased with body mass (Figure 2). Ordinary least squares regression indicated a positive association (slope = 0.096LJ°CLJgLJ¹), with very high explained variance (R² = 0.98). Bootstrapping (10,000 replicates) confirmed the robustness of this slope estimate, yielding a 95% confidence interval of 0.064-0.149LJ°CLJgLJ¹, indicating relatively high precision. This slope implies that a 10 g increase in body mass corresponds to an approximately 1 °C higher HTL. Permutation testing further supported the significance of this relationship (two-sided p = 0.017). In addition, Bayesian regression produced a posterior median slope of 0.094 (95% credible interval: 0.055–0.146), fully consistent with the frequentist analyses.

**Figure 2.**
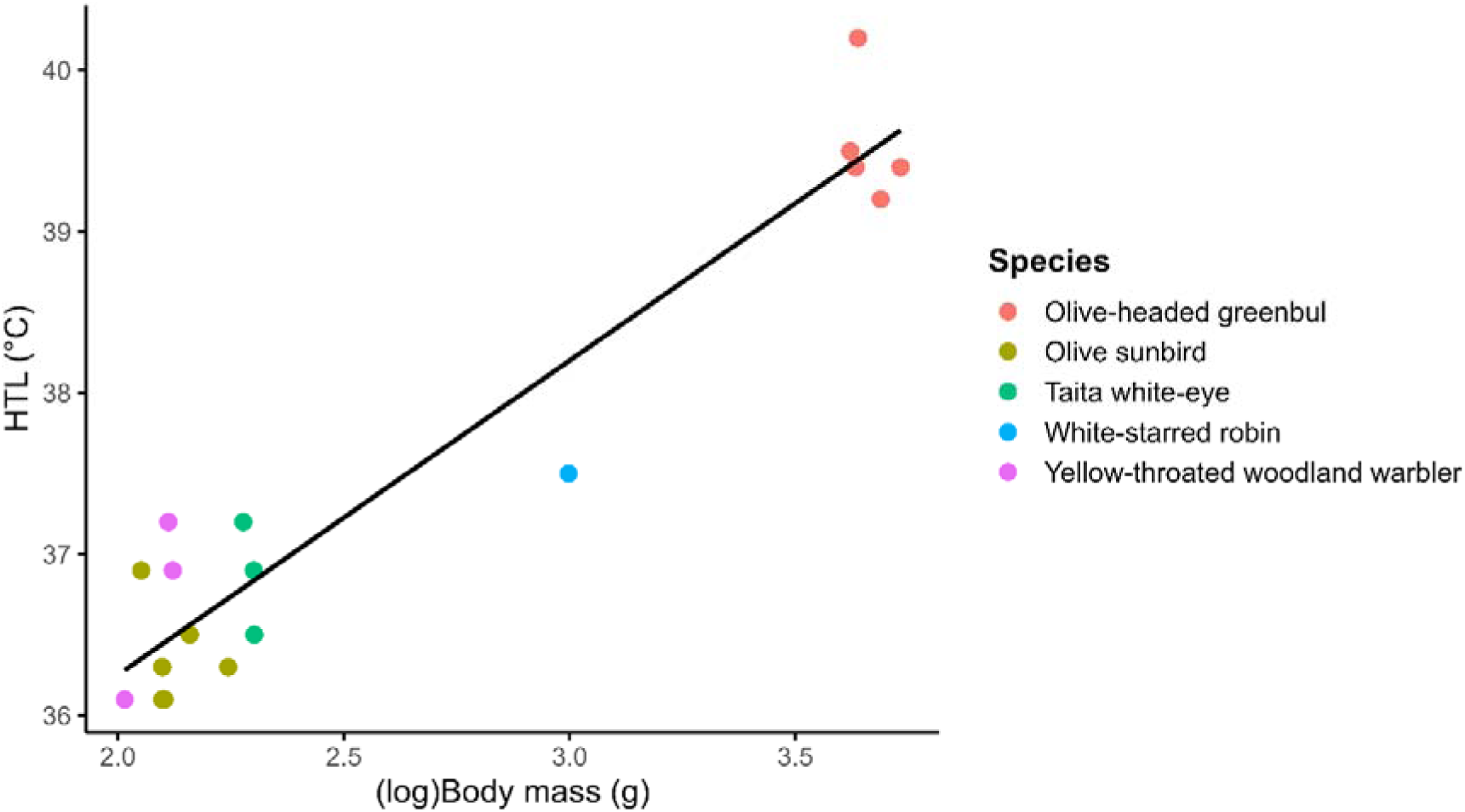
Relationship between average log-transformed body mass (g) and heat tolerance limit (HTL. °C) across five bird species. Points represent individual birds. coloured by species. and black lines show the linear regression fit for all data.

Inflection points for evaporative cooling capacity EHL/MHP varied among species, ranging from 25LJ°C in the olive sunbird and yellow-throated woodland warbler to 32LJ°C in the olive-headed greenbul (Table S2 – Supplementary material). Slopes above the inflection temperature differed in both magnitude and direction. The olive sunbird and Taita white-eye showed very small negative slopes (−0.0024 and −0.0056, respectively), effectively indistinguishable from zero, whereas the olive-headed greenbul and white-starred robin exhibited substantially positive slopes (0.0313 and 0.0443, respectively). These slopes reflect how rapidly evaporative heat loss increased with temperature once the inflection point was reached, with steeper positive values indicating a stronger capacity to enhance evaporative cooling at higher air temperatures. Linear regression of slope against mean body mass revealed a positive, but non-significant, trend (slope = 0.00124, SE = 0.00074, t = 1.68, p = 0.19, R² = 0.49; Figure 3). This indicates that as expected (prediction 2b) larger species tended to increase evaporative cooling more steeply with temperature. Overall, none of the species approached an EHL/MHP ratio of 1 (ratio ranged from 0.31 to 0.51, Table 2) highlighting that evaporative cooling efficiency was ‘weak’ across all species (prediction 3).

**Figure 3.**
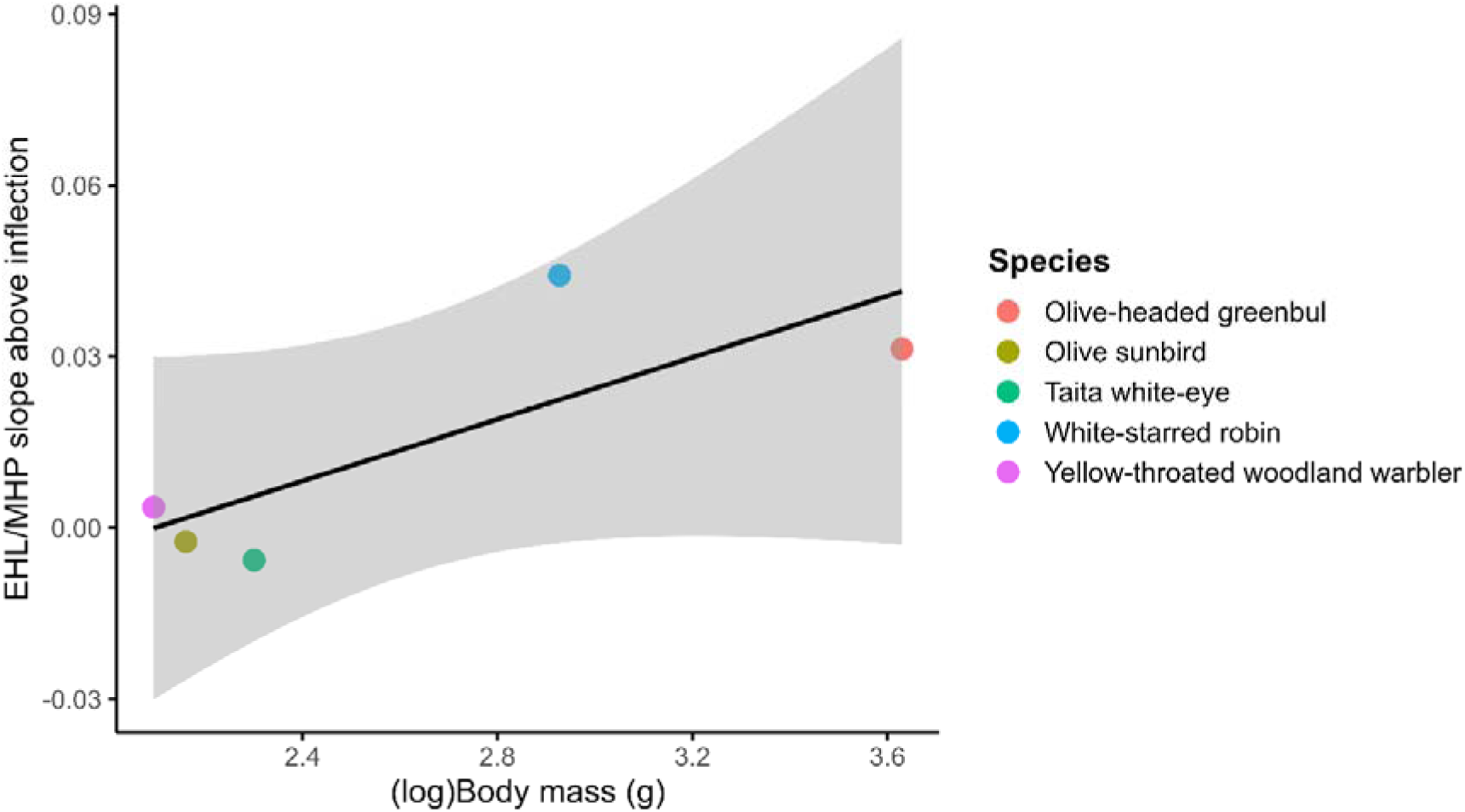
Relationship between species average log-transformed body mass (g) and evaporative cooling efficiency (EHL/MHP slope above inflection) across five bird species. Points are coloured by species. and the black line represents the linear regression across all individuals.

**Table 2.**
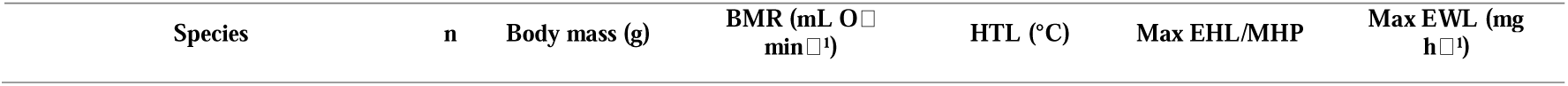
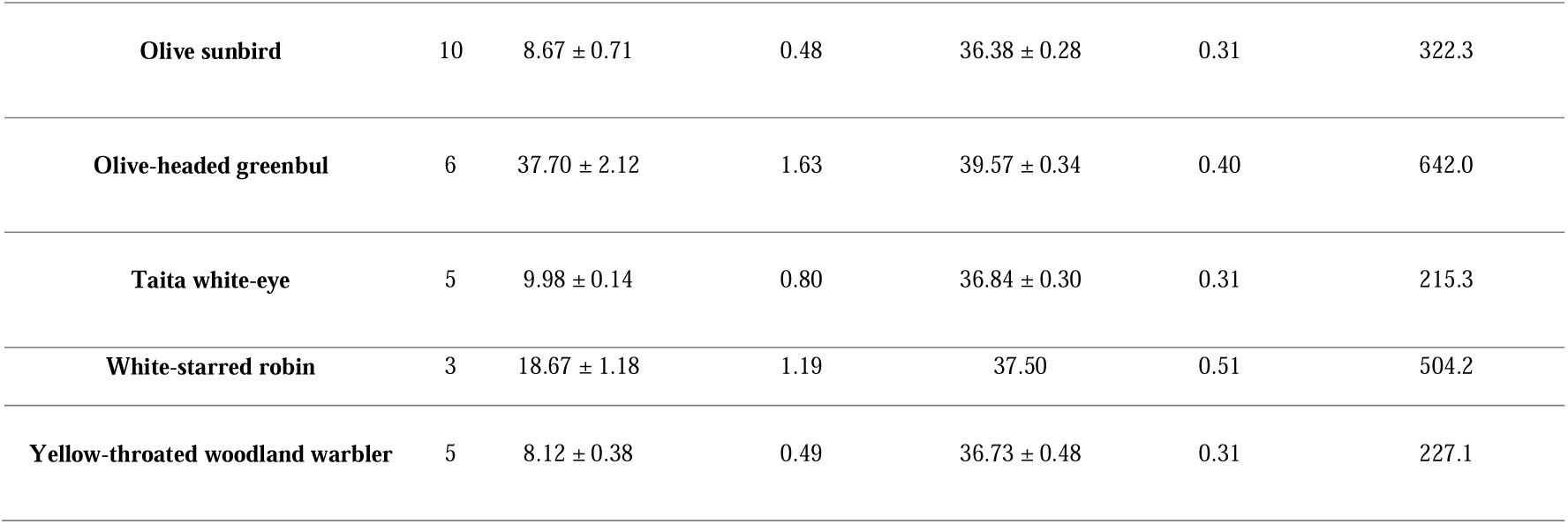
Values (±LJs.d.. where applicable) of average body mass (g) and physiological thermoregulatory traits measured in five forest-dependent bird species from the Taita Hills. Kenya. BMR (mL OLJ minLJ¹) is the basal metabolic rate. estimated as the lowest observed resting metabolic rate (RMR); HTL (°C) is the mean heat tolerance limit; Max EHL/MHP is the maximum ratio of evaporative heat loss to metabolic heat production; EWL (mg hLJ¹) is the maximum evaporative water loss recorded.

| Species | n | Body mass (g) | BMR (mL O<sub>2</sub> min<sup>-1</sup>) | HTL (°C) | Max EHL/MHP | Max EWL (mg h<sup>-1</sup>) |
| --- | --- | --- | --- | --- | --- | --- |
| **Olive sunbird** | 10 | 8.67 ± 0.71 | 0.48 | 36.38 ± 0.28 | 0.31 | 322.3 |
| **Olive-headed greenbul** | 6 | 37.70 ± 2.12 | 1.63 | 39.57 ± 0.34 | 0.40 | 642.0 |
| **Taita white-eye** | 5 | 9.98 ± 0.14 | 0.80 | 36.84 ± 0.30 | 0.31 | 215.3 |
| **White-starred robin** | 3 | 18.67 ± 1.18 | 1.19 | 37.50 | 0.51 | 504.2 |
| **Yellow-throated woodland warbler** | 5 | 8.12 ± 0.38 | 0.49 | 36.73 ± 0.48 | 0.31 | 227.1 |

Finally, to place our findings in a broader biogeographic context (prediction 4), we compared our estimated critical temperatures of the Taita Hills bird species with the global dataset of Khaliq et al. (2015). Because the metabolic responses of all our study species were predominantly V-shaped, lower and upper critical temperatures effectively coincided, and for this comparison we therefore treated them as equivalent. When compared with the global dataset, the Taita Hills species consistently fell near the lower end of the distributions for critical temperatures. This pattern was even more pronounced when restricting comparisons to tropical and subtropical species (latitude ±30°; n = 24), with most study species below the 10th percentile for both LCT and UCT (Figure 4). The elevation of the Taita Hills study site (∼1,800LJm) is higher than the maximum elevation represented in the tropical small-bird subset of the global dataset (max = 1,514LJm), indicating that the montane conditions of our study site are above those captured by the available comparative data.

**Figure 4.**
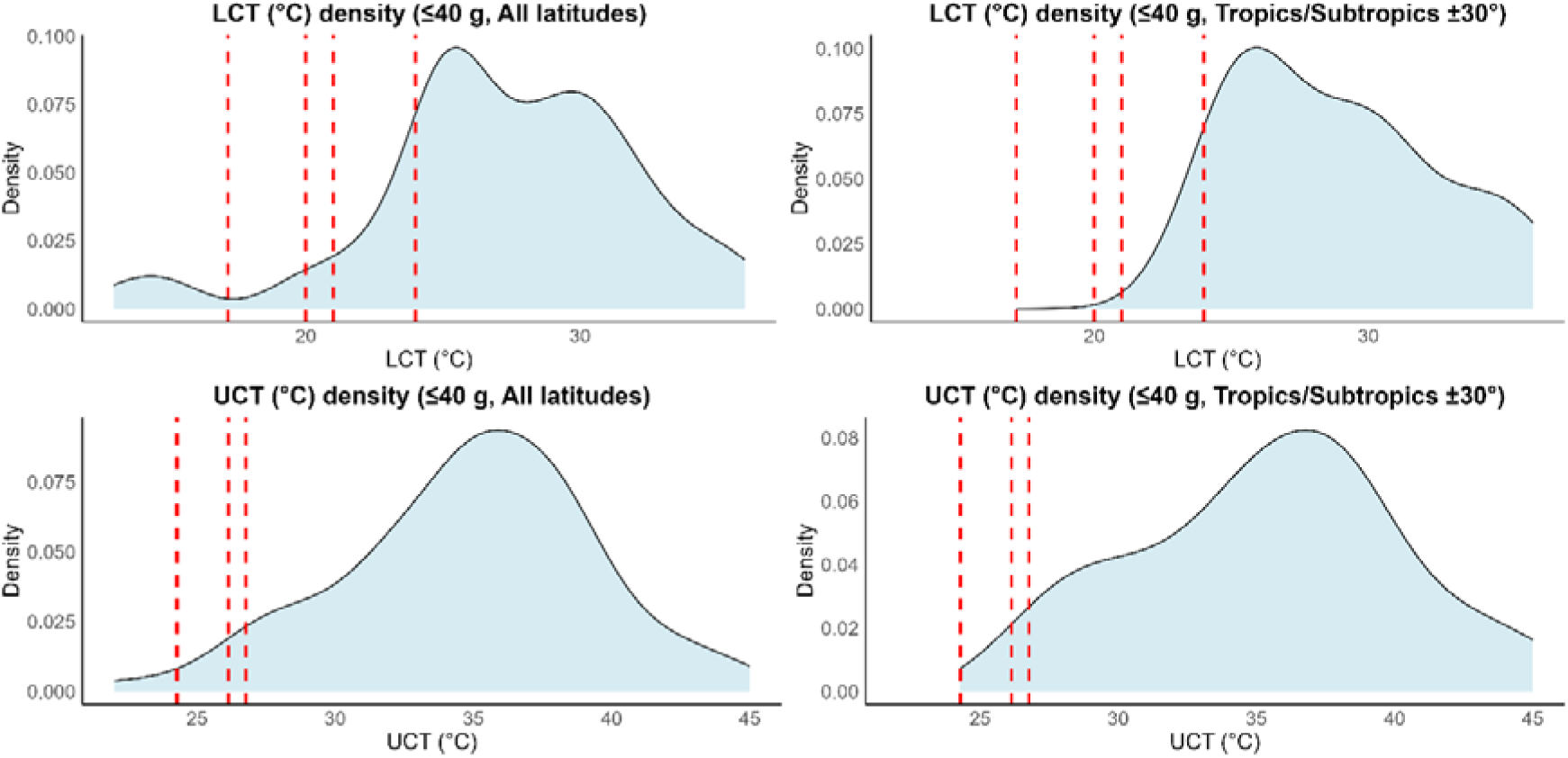
Density distributions of lower (LCT) and upper (UCT) critical temperatures for small bird species (≤ 40 g) based on data from Khaliq et al. (2015). Distributions are shown for all latitudes (left panels) and for tropical/subtropical species within ±30° latitude (right panels). Vertical dashed lines indicate values measured in this study (apart from the white-starred robin; see above). Note: in the current analysis. dashed lines correspond to breakpoints from segmented models representing LCT and UCT. In most species. these breakpoints occur close together. reflecting a narrow or weakly defined thermoneutral zone. which in previous V-shaped fits was represented by a single minimum.

## Discussion

This study assessed the thermoregulatory capacities of five tropical forest-dependent songbirds from the fragmented montane forests of the Taita Hills. Contrary to expectation, the relationship between resting metabolic rate (RMR) and ambient temperature was predominantly V-shaped, suggesting the absence, or at best the existence of a very narrow, TNZ. In line with prediction 2a, larger species showed greater heat tolerance: HTLs increased with body mass, with all analyses supporting a robust positive relationship. Prediction 2b, that larger species would show higher evaporative cooling capacity, was partly supported. Of the five study species, only the olive-headed greenbul and the white-starred robin, the two largest species in our sample, showed evidence of increasing evaporative heat loss with rising temperature, indicating some capacity to enhance thermoregulatory performance under warmer conditions. In contrast, the smaller species showed no detectable change in evaporative cooling with increasing temperature. As expected under prediction 3, evaporative cooling efficiency remained low across all species (EHL/MHPLJ<LJ1), suggesting that tropical montane birds operate with little physiological margin to dissipate metabolic heat. Finally, consistent with prediction 4, the lower and upper critical temperatures of the Taita Hills species were shifted toward the lower end of the global distributions for small-bodied birds (Khaliq et al., 2015), particularly when comparisons were restricted to tropical and subtropical species. The combination of narrow thermal ranges, low critical temperatures, and limited capacity or flexibility in thermoregulatory capacity indicates that montane forest birds, particularly the smaller species, may be especially vulnerable to climate change–induced increases in ambient temperature.

Recent work has questioned the long-standing assumption that tropical birds universally possess narrow thermoneutral zones and low flexibility in thermoregulation. Monge et al. (2023), for example, argue that thermal physiology among tropical birds is more variable than previously thought, with some species maintaining body temperature across a broader range of ambient conditions. Our results, however, align more closely with the traditional view, as the Taita Hills species appear restricted to low critical temperatures and exhibit limited thermoregulatory flexibility (Figure 1-3), traits consistent with specialization to the cool, stable microclimates of tropical montane forests. The relatively high elevation of our study site (∼1,800 m) further suggests that our study species may be adapted to conditions cooler than those experienced by most tropical small-bodied species. Consistent with this interpretation, HTLs were also relatively low compared with passerines from southern Africa and the American Southwest (McKechnie et al., 2017; Smith et al., 2017; Whitfield et al., 2015), reinforcing the view that these montane birds remain within their thermoneutral range only across a narrow band of ambient temperatures and must expend additional metabolic energy to regulate body temperature once environmental conditions deviate from this range. Nevertheless, we cannot exclude the possibility that our results reflect a seasonal snapshot rather than fixed physiological limits. Seasonal plasticity in metabolic rates, as discussed by Monge et al. (2023), could allow broader thermal tolerance under different climatic conditions. It is thus important to note that our measurements were conducted during the cool–wet season only. Khaliq et al.’s (2015) data for small birds indicate LCT values that can vary seasonally, typically ∼28LJ°C in the warm season and ∼24LJ°C in the cool season, the latter closely matching the mean temperature of minimum metabolic rate across our five study species (Figure 4). The relatively low thermal limits observed in our study may partly reflect the seasonal context of our measurements.

Body mass influenced thermal physiology across species, with larger birds tolerating higher ambient temperatures than smaller species (Table 2, Figure 2-3). According to Bergmann’s rule, smaller body sizes are generally favoured in warmer climates because a higher surface-area-to-volume ratio enhances heat dissipation (Bergmann, 1948). However, this same property also causes smaller individuals to exchange heat with their environment more rapidly, leading to faster gains as well as losses of heat. In contrast, larger birds, with lower surface-area-to-volume ratios, possess greater thermal inertia: they warm up more slowly and thus experience delayed increases in body temperature during acute heat exposure (Conradie et al., 2020; McNab, 2002). Moreover, smaller species have proportionally smaller body water reserves and higher mass-specific rates of evaporative cooling, which can accelerate dehydration and further limit their capacity to tolerate heat (McKechnie et al., 2021). Together, these mechanisms likely explain why, in the short-term physiological context of our study, individuals of larger species endured higher ambient temperatures before reaching their thermal limits. Evaporative cooling capacity, however, showed little relationship with body size and remained low across all species (all ratios < 0.51, Table 2), contrasting with arid-zone passerines that can dissipate more than twice their metabolic heat production through evaporation (McKechnie et al., 2016). Similarly, comparative data indicate that lowland tropical birds achieve greater evaporative efficiency and tolerate hyperthermia more effectively than montane species (Cabello-Vergel et al., 2024), suggesting that cool, humid montane environments favour reduced reliance on evaporative cooling (Smith et al., 2017; Thompson et al., 2018; van de Ven et al., 2019). Taken together, our results indicate that all species studied here likely have a physiological profile consistent with adaptation to mild, thermally stable environments but with limited capacity to dissipate excess heat.

By characterizing thermoregulatory limits in tropical montane birds inhabiting the Kenyan Taita Hills area, this study contributes to a better understanding of how forest-dependent species cope with heat stress, although it was based on relatively small sample sizes and measurements from a single season and site. Broader sampling across seasons, years and sites would help confirm the robustness of the observed patterns. In addition, HTLs were measured in dry rather than humidified air. Although this approach follows established protocols and facilitates comparison across studies, it likely overestimates the heat tolerance that birds experience in the naturally humid montane conditions of the Taita Hills. Despite these limitations, our findings indicate that in our study area, small tropical montane forest birds likely operate within narrow thermal margins and may have limited evaporative cooling capacity, suggesting high sensitivity to heat stress. Warming trends and increasingly frequent heat waves across Africa (Amou et al., 2021; Nying’uro et al., 2024), together with microclimatic shifts from forest fragmentation or edge exposure, could therefore elevate energetic and reproductive costs, placing these communities at risk under continued climate change. Future work combining more extensive empirical data across seasons, years, and individuals with predictive models of microclimate and habitat change will be crucial to anticipate how tropical montane birds respond to a rapidly warming and increasingly fragmented world.

## Acknowledgements

The authors would like to thank Laurence Chovu, John Maghanga, Musa Makomba, Nathaniel Mkombola, Nathaniel Ndighila, Caoimhe Abdul-Wahab, and Lauren Brickle for their help during data collection.

## Author contributions

Conceptualization: C.P., R.X., B.A., D.S., L.L.; Methodology: C.P., R.X., D.S.; Fieldwork and data collection: C.P., B.A. Data analysis: C.P., R.X. Visualization: C.P., R.X. Funding acquisition: B.A., L.L.; Administrative and logistical support: P.N., M.G.; Project supervision: B.A., F.V., D.S., L.L. Writing – original draft: C.P, R.X.; Writing – review & editing: B.A., F.V., P.N., M.G., D.S., L.L.

## Data availability

The data and the R script supporting the findings of this study will be made publicly available in an open-access repository upon publication.

## References

1. Albright, T. P., Mutiibwa, D., Gerson, Alexander R., Smith, E. K., Talbot, W. A., O’Neill, J. J., McKechnie, A. E., & Wolf, B. O. (2017). Mapping evaporative water loss in desert passerines reveals an expanding threat of lethal dehydration. Proceedings of the National Academy of Sciences, 114(9), 2283–2288. 10.1073/pnas.1613625114

2. Alroy, J. (2017). Effects of habitat disturbance on tropical forest biodiversity. Proceedings of the National Academy of Sciences, 114(23), 6056–6061. 10.1073/pnas.1611855114

3. Amou, M., Gyilbag, A., Demelash, T., & Xu, Y. (2021). Heatwaves in Kenya 1987–2016: Facts from CHIRTS High Resolution Satellite Remotely Sensed and Station Blended Temperature Dataset. Atmosphere, 12(1), 37. 10.3390/atmos12010037

4. Araújo, M. B., Ferri-Yáñez, F., Bozinovic, F., Marquet, P. A., Valladares, F., & Chown, S. L. (2013). Heat freezes niche evolution. Ecology Letters, 16(9), 1206–1219. 10.1111/ele.12155

5. Barlow, J., Lennox, G. D., Ferreira, J., Berenguer, E., Lees, A. C., Nally, R. M., Thomson, J. R., Ferraz, S. F. de B., Louzada, J., Oliveira, V. H. F., Parry, L., Ribeiro de Castro Solar, R., Vieira, I. C. G., Aragão, L. E. O. C., Begotti, R. A., Braga, R. F., Cardoso, T. M., de Oliveira, R. C., Souza Jr, C. M., … Gardner, T. A. (2016). Anthropogenic disturbance in tropical forests can double biodiversity loss from deforestation. Nature, 535(7610), Article 7610. 10.1038/nature18326

6. Bozinovic, F., & Pörtner, H.-O. (2015). Physiological ecology meets climate change. Ecology and Evolution, 5(5), 1025–1030. 10.1002/ece3.1403

7. Burgess, N. D., Butynski, T. M., Cordeiro, N. J., Doggart, N. H., Fjeldså, J., Howell, K. M., Kilahama, F. B., Loader, S. P., Lovett, J. C., Mbilinyi, B., Menegon, M., Moyer, D. C., Nashanda, E., Perkin, A., Rovero, F., Stanley, W. T., & Stuart, S. N. (2007). The biological importance of the Eastern Arc Mountains of Tanzania and Kenya. Biological Conservation, 134(2), 209–231. 10.1016/j.biocon.2006.08.015

8. Cabello-Vergel, J., González-Medina, E., Parejo, M., Abad-Gómez, J. M., Playà-Montmany, N., Patón, D., Sánchez-Guzmán, J. M., Masero, J. A., Gutiérrez, J. S., & Villegas, A. (2022). Heat tolerance limits of Mediterranean songbirds and their current and future vulnerabilities to temperature extremes. Journal of Experimental Biology, 225(23), jeb244848. 10.1242/jeb.244848

9. Cabello-Vergel, J., Gutiérrez, J. S., González-Medina, E., Sánchez-Guzmán, J. M., Masero, J. A., & Villegas, A. (2024). Seasonal and between-population variation in heat tolerance and cooling efficiency in a Mediterranean songbird. Journal of Thermal Biology, 125, 103977. 10.1016/j.jtherbio.2024.103977

10. Carroll, J. M., Davis, C. A., Elmore, R. D., Fuhlendorf, S. D., & Thacker, E. T. (2015). Thermal patterns constrain diurnal behavior of a ground-dwelling bird. Ecosphere, 6(11), art222. 10.1890/ES15-00163.1

11. Conradie, S. R., Woodborne, S. M., Wolf, B. O., Pessato, A., Mariette, M. M., & McKechnie, A. E. (2020). Avian mortality risk during heat waves will increase greatly in arid Australia during the 21st century. Conservation Physiology, 8(1), coaa048. 10.1093/conphys/coaa048

12. Haddad, N. M., Brudvig, L. A., Clobert, J., Davies, K. F., Gonzalez, A., Holt, R. D., Lovejoy, T. E., Sexton, J. O., Austin, M. P., Collins, C. D., Cook, W. M., Damschen, E. I., Ewers, R. M., Foster, B. L., Jenkins, C. N., King, A. J., Laurance, W. F., Levey, D. J., Margules, C. R., … Townshend, J. R. (2015). Habitat fragmentation and its lasting impact on Earth’s ecosystems. Science Advances, 1(2), e1500052. 10.1126/sciadv.1500052

13. IPCC. (2023). Climate Change 2022 – Impacts, Adaptation and Vulnerability: Working Group II Contribution to the Sixth Assessment Report of the Intergovernmental Panel on Climate Change (1st ed.). Cambridge University Press. 10.1017/9781009325844

14. Keinath, D. A., Doak, D. F., Hodges, K. E., Prugh, L. R., Fagan, W., Sekercioglu, C. H., Buchart, S. H. M., & Kauffman, M. (2017). A global analysis of traits predicting species sensitivity to habitat fragmentation. Global Ecology and Biogeography, 26(1), 115–127. 10.1111/geb.12509

15. Lighton, J. R. B. (2018). Measuring Metabolic Rates: A Manual for Scientists (2nd ed.). Oxford University Press. 10.1093/oso/9780198830399.001.0001

16. Linke, J. (2007). Introduction: Structure, function and change of forest landscapes. https://www.academia.edu/17511717/Introduction_structure_function_and_change_of_forest_landscapes

17. Lovett, J. C., & Wasser, S. K. (2008). Biogeography and ecology of the rain forests of eastern Africa. Biogeography and Ecology of the Rain Forests of Eastern Africa. https://www.cabdirect.org/cabdirect/abstract/20093194721

18. Marcondes, R. S. (2019). Realistic scenarios of missing taxa in phylogenetic comparative methods and their effects on model selection and parameter estimation. PeerJ, 7, e7917. 10.7717/peerj.7917

19. McKechnie, A. E., Gerson, A. R., McWhorter, T. J., Smith, E. K., Talbot, W. A., & Wolf, B. O. (2017). Avian thermoregulation in the heat: Evaporative cooling in five Australian passerines reveals within-order biogeographic variation in heat tolerance. Journal of Experimental Biology, 220(13), 2436–2444. 10.1242/jeb.155507

20. McKechnie, A. E., Gerson, A. R., & Wolf, B. O. (2021). Thermoregulation in desert birds: Scaling and phylogenetic variation in heat tolerance and evaporative cooling. Journal of Experimental Biology, 224(Suppl_1), jeb229211. 10.1242/jeb.229211

21. McKechnie, A. E., Rushworth, I. A., Myburgh, F., & Cunningham, S. J. (2021). Mortality among birds and bats during an extreme heat event in eastern South Africa. Austral Ecology, 46(4), 687–691. 10.1111/aec.13025

22. Mckechnie, A. E., & Swanson, D. L. (2010). Sources and significance of variation in basal, summit and maximal metabolic rates in birds. Current Zoology, 56(6), 741–758. 10.1093/czoolo/56.6.741

23. McKechnie, A. E., Whitfield, M. C., Smit, B., Gerson, A. R., Smith, E. K., Talbot, W. A., McWhorter, T. J., & Wolf, B. O. (2016). Avian thermoregulation in the heat: Efficient evaporative cooling allows for extreme heat tolerance in four southern hemisphere columbids. Journal of Experimental Biology, 219(14), 2145–2155. 10.1242/jeb.138776

24. McKechnie, A. E., & Wolf, B. O. (2009). Climate change increases the likelihood of catastrophic avian mortality events during extreme heat waves. Biology Letters, 6(2), 253–256. 10.1098/rsbl.2009.0702

25. McNab, B. K. (2002). The Physiological Ecology of Vertebrates: A View from Energetics. Cornell University Press.

26. McNab, B. K. (2012). Extreme Measures: The Ecological Energetics of Birds and Mammals. University of Chicago Press.

27. Monge, O., Maggini, I., Schulze, C. H., Dullinger, S., & Fusani, L. (2023). Physiologically vulnerable or resilient? Tropical birds, global warming, and redistributions. Ecology and Evolution, 13(4), e9985. 10.1002/ece3.9985

28. Mulwa, M., Teucher, M., Ulrich, W., & Habel, J. C. (2021). Bird communities in a degraded forest biodiversity hotspot of East Africa. Biodiversity and Conservation, 30(8), 2305–2318. 10.1007/s10531-021-02190-y

29. Mundry, R. (2014). Statistical Issues and Assumptions of Phylogenetic Generalized Least Squares. In L. Z. Garamszegi (Ed.), Modern Phylogenetic Comparative Methods and Their Application in Evolutionary Biology (pp. 131–153). Springer Berlin Heidelberg. 10.1007/978-3-662-43550-2_6

30. Myers, N., Mittermeier, R. A., Mittermeier, C. G., da Fonseca, G. A. B., & Kent, J. (2000). Biodiversity hotspots for conservation priorities. Nature, 403(6772), Article 6772. 10.1038/35002501

31. Newmark, W. D. (2002). Conserving Biodiversity in East African Forests: A Study of the Eastern Arc Mountains. Springer Science & Business Media.

32. O’Connor, R. S., Le Pogam, A., Young, K. G., Robitaille, F., Choy, E. S., Love, O. P., Elliott, K. H., Hargreaves, A. L., Berteaux, D., Tam, A., & Vézina, F. (2021). Limited heat tolerance in an Arctic passerine: Thermoregulatory implications for coldLJspecialized birds in a rapidly warming world. Ecology and Evolution, 11(4), 1609–1619. 10.1002/ece3.7141

33. Osváth, G., Daubner, T., Dyke, G., Fuisz, T. I., Nord, A., Pénzes, J., Vargancsik, D., Vágási, C. I., Vincze, O., & Pap, P. L. (2018). How feathered are birds? Environment predicts both the mass and density of body feathers. Functional Ecology, 32(3), 701–712. 10.1111/1365-2435.13019

34. Pearse, W. D., Davies, T. J., & Wolkovich, E. M. (2025). How to Define, Use, and Interpret Pagel’s \ łambda \ (Lambda) in Ecology and Evolution. Global Ecology and Biogeography, 34(4), e70012. 10.1111/geb.70012

35. Pellikka, P. K. E., Clark, B. J. F., Gosa, A. G., Himberg, N., Hurskainen, P., Maeda, E., Mwang’ombe, J., Omoro, L. M. A., & Siljander, M. (2013). Chapter 13—Agricultural Expansion and Its Consequences in the Taita Hills, Kenya. In P. Paron, D. O. Olago, & C. T. Omuto (Eds.), Developments in Earth Surface Processes (Vol. 16, pp. 165–179). Elsevier. 10.1016/B978-0-444-59559-1.00013-X

36. Pellikka, P. K. E., Lötjönen, M., Siljander, M., & Lens, L. (2009). Airborne remote sensing of spatiotemporal change (1955–2004) in indigenous and exotic forest cover in the Taita Hills, Kenya. International Journal of Applied Earth Observation and Geoinformation, 11(4), 221–232. 10.1016/j.jag.2009.02.002

37. Polato, N. R., Gill, B. A., Shah, A. A., Gray, M. M., Casner, K. L., Barthelet, A., Messer, P. W., Simmons, M. P., Guayasamin, J. M., Encalada, A. C., Kondratieff, B. C., Flecker, A. S., Thomas, S. A., Ghalambor, C. K., Poff, N. L., Funk, W. C., & Zamudio, K. R. (2018). Narrow thermal tolerance and low dispersal drive higher speciation in tropical mountains. Proceedings of the National Academy of Sciences of the United States of America, 115(49), 12471–12476. 10.1073/pnas.1809326115

38. Pollock, H. S., Brawn, J. D., & Cheviron, Z. A. (2021). Heat tolerances of temperate and tropical birds and their implications for susceptibility to climate warming. Functional Ecology, 35(1), 93–104. 10.1111/1365-2435.13693

39. Rheindt, F. E., Donald, P. F., Donsker, D. B., Gerbracht, J. A., Iliff, M. J., Lepage, D., Norman, J. A., Rasmussen, P. C., Schodde, R., Schulenberg, T. S., Areta, J. I., Brammer, F. P., Chesser, R. T., Dowsett, R. J., Peterson, A., Alström, P., Stervander, M., Remsen, J. V., Garnett, S. T., … Christidis, L. (2025). AviList: A unified global bird checklist. Biodiversity and Conservation, 34(10), 3359–3376. 10.1007/s10531-025-03120-y

40. Rogalla, S., Patil, A., Dhinojwala, A., Shawkey, M. D., & D’Alba, L. (2021). Enhanced photothermal absorption in iridescent feathers. Journal of the Royal Society, Interface, 18(181), 20210252. 10.1098/rsif.2021.0252

41. Ruuskanen, S., Hsu, B.-Y., & Nord, A. (2021). Endocrinology of thermoregulation in birds in a changing climate. Molecular and Cellular Endocrinology, 519, 111088. 10.1016/j.mce.2020.111088

42. Scheffers, B. R., Evans, T. A., Williams, S. E., & Edwards, D. P. (2014). Microhabitats in the tropics buffer temperature in a globally coherent manner. Biology Letters, 10(12), 20140819. 10.1098/rsbl.2014.0819

43. Schmidt-Nielsen, K. (1997). Animal Physiology: Adaptation and Environment. Cambridge University Press.

44. Schwenk, K., Padilla, D. K., Bakken, G. S., & Full, R. J. (2009). Grand challenges in organismal biology. Integrative and Comparative Biology, 49(1), 7–14. 10.1093/icb/icp034

45. Smith, E. K., O’Neill, J. J., Gerson, A. R., McKechnie, A. E., & Wolf, B. O. (2017). Avian thermoregulation in the heat: Resting metabolism, evaporative cooling, and heat tolerance in Sonoran Desert songbirds. Journal of Experimental Biology, jeb.161141. 10.1242/jeb.161141

46. Stillman, J. H. (2019). Heat Waves, the New Normal: Summertime Temperature Extremes Will Impact Animals, Ecosystems, and Human Communities. Physiology, 34(2), 86–100. 10.1152/physiol.00040.2018

47. Swanson, D. L. (2010). Seasonal Metabolic Variation in Birds: Functional and Mechanistic Correlates. In C. F. Thompson (Ed.), Current Ornithology Volume 17 (pp. 75–129). Springer. 10.1007/978-1-4419-6421-2_3

48. Taubert, F., Fischer, R., Groeneveld, J., Lehmann, S., Müller, M. S., Rödig, E., Wiegand, T., & Huth, A. (2018). Global patterns of tropical forest fragmentation. Nature, 554(7693), Article 7693. 10.1038/nature25508

49. Terschanski, J., Nunes, M. H., Aalto, I., Pellikka, P., Wekesa, C., & Maeda, E. E. (2024). The role of vegetation structural diversity in regulating the microclimate of human-modified tropical ecosystems. Journal of Environmental Management, 360, 121128. 10.1016/j.jenvman.2024.121128

50. Teucher, M., Schmitt, C. B., Wiese, A., Apfelbeck, B., Maghenda, M., Pellikka, P., Lens, L., & Habel, J. C. (2020). Behind the fog: Forest degradation despite logging bans in an East African cloud forest. Global Ecology and Conservation, 22, e01024. 10.1016/j.gecco.2020.e01024

51. Thompson, M. L., Cunningham, S. J., & McKechnie, A. E. (2018). Interspecific variation in avian thermoregulatory patterns and heat dissipation behaviours in a subtropical desert. Physiology & Behavior, 188, 311–323. 10.1016/j.physbeh.2018.02.029

52. Tracy, R. (2010). *Properties of Air* 2010.

53. Tuff, K. T., Tuff, T., & Davies, K. F. (2016). A framework for integrating thermal biology into fragmentation research. Ecology Letters, 19(4), 361–374. 10.1111/ele.12579

54. van de Ven, T. M. F. N., McKechnie, A. E., & Cunningham, S. J. (2019). The costs of keeping cool: Behavioural trade-offs between foraging and thermoregulation are associated with significant mass losses in an arid-zone bird. Oecologia, 191(1), 205–215. 10.1007/s00442-019-04486-x

55. van de Ven, T. M. F. N., Mzilikazi, N., & McKechnie, A. E. (2013). Seasonal Metabolic Variation in Two Populations of an Afrotropical Euplectid Bird. Physiological and Biochemical Zoology, 86(1), 19–26. 10.1086/667989

56. Whitfield, M. C., Smit, B., McKechnie, A. E., & Wolf, B. O. (2015). Avian thermoregulation in the heat: Scaling of heat tolerance and evaporative cooling capacity in three southern African arid-zone passerines. Journal of Experimental Biology, 218(11), 1705–1714. 10.1242/jeb.121749

57. Wolf, B. O., & Walsberg, G. E. (1996). Thermal Effects of Radiation and Wind on a Small Bird and Implications for Microsite Selection. Ecology, 77(7), 2228–2236. 10.2307/2265716

