## Supplementary material for "Weak evaporative cooling capacity and body size shape thermal limits in tropical montane forest birds"

| Species | T_min (°C) | SE |
| --- | --- | --- |
| Olive sunbird | 24.91 | 0.43 |
| Olive-headed greenbul | 26.23 | 0.25 |
| Taita white-eye | 23.72 | 0.37 |
| White-starred robin | 25.04 | 0.69 |
| Yellow-throated woodland warbler | 24.08 | 0.54 |

Table S1. Values represent the temperatures (T_min (°C)) at which minimum resting metabolic rate (RMR) was achieved. with associated standard errors (SE) derived from species-specific segmented regressions.

| Species | Inflection point (°C) | Slope above inflection | Slope SE | t value |
| --- | --- | --- | --- | --- |
| Olive sunbird | 28.00 | -0.002 | 0.011 | -0.21 |
| Olive-headed greenbul | 32.00 | 0.031 | 0.019 | 1.66 |
| Taita white-eye | 28.62 | -0.006 | 0.003 | -2.13 |
| White-starred robin | 28.59 | 0.044 | 0.013 | 3.42 |
| Yellow-throated woodland warbler | 25.00 | 0.004 | 0.004 | 0.98 |

Table S2. Results from segmented regression analyses of evaporative cooling efficiency (EHL/MHP) versus ambient temperature. Inflection points (°C) represent estimated ambient temperatures at which the slope of EHL/MHP changes. Slope above inflection. standard error (Slope SE). and t value refer to the linear relationship of EHL/MHP with temperature for values above the inflection point.
